# Characterizing RNA structures *in vitro* and *in vivo* with selective 2’-hydroxyl acylation analyzed by primer extension sequencing (SHAPE-Seq)

**DOI:** 10.1101/034470

**Authors:** Kyle E. Watters, Angela M Yu, Eric J. Strobel, Alex H. Settle, Julius B. Lucks

**Affiliations:** School of Chemical and Biomolecular Engineering, Cornell University, Ithaca NY 14853; Tri-Institutional Program in Computational Biology and Medicine, Cornell University, Ithaca, New York, Weill Cornell Medical College, New York, New York, Memorial Sloan-Kettering Cancer Center, New York, New York; Computational Biology Program, Memorial Sloan-Kettering Cancer Center, New York, New York 10065

**Keywords:** RNA, SHAPE, SHAPE-Seq, RNA Structure Probing, RNA Structure Prediction, *in vivo* RNA structure, RNA-Ligand Interactions

## Abstract

RNA molecules adopt a wide variety of structures that perform many cellular functions, including catalysis, small molecule sensing, and cellular defense, among others. Our ability to characterize, predict, and design RNA structures are key factors for understanding and controlling the biological roles of RNAs. Fortunately, there has been rapid progress in this area, especially with respect to experimental methods that can characterize RNA structures in a high throughput fashion using chemical probing and next-generation sequencing. Here, we describe one such method, selective 2’-hydroxyl acylation analyzed by primer extension sequencing (SHAPE-Seq), which measures nucleotide resolution flexibility information for RNAs *in vitro* and *in vivo*. We outline the process of designing and performing a SHAPE-Seq experiment and describe methods for using experimental SHAPE-Seq data to restrain computational folding algorithms to generate more accurate predictions of RNA secondary structure. We also provide a number of examples of SHAPE-Seq reactivity spectra obtained *in vitro* and *in vivo* and discuss important considerations for performing SHAPE-Seq experiments, both in terms of collecting and analyzing data. Finally we discuss improvements and extensions of these experimental and computational techniques that promise to deepen our knowledge of RNA folding and function.

## 1. Introduction

The dual informational/structural nature of RNA molecules allows them to simultaneously encode genetic information and actively direct cellular processes. Many RNAs assume highly sophisticated structures that mediate a diverse set of functions. These functions range from catalysis, as in the case of RNA enzymes like RNase P [1] and the ribosome, to a diverse and expanding array of regulatory mechanisms including riboswitches [2,3], RNAi [4], RNA transcriptional attenuators [5], CRISPR [6], thermometers [7], and many others. A large number of these RNAs are non-coding (ncRNA) and function in a purely structural manner without carrying genetic information [8-10]. Our understanding of the importance of these functional ncRNAs is increasing and many more continue to be discovered at a rapid pace [10]. Thus, the development of tools to quickly and accurately characterize the structure-function relationships of ncRNAs is essential to advancing the field of RNA biology [11].

A common method of characterizing RNA structure is to isolate the RNA of interest *in vitro* and perform enzymatic or chemical probing experiments that reveal information about an RNA molecule’s secondary and tertiary structure [12,13]. These experiments interrogate RNA structures by measuring nucleotide accessibility for RNase cleavage or chemical modification and can be used to infer whether a nucleotide within an RNA molecule is predominantly single- or double-stranded [12]. Chemical probes have become more frequently used due to their higher resolution and ability to transport across membranes to react with RNAs inside cells. These probes use a range of chemistries to covalently modify RNAs is a structure-dependent fashion and can be roughly divided into three classes [12]: 1) base-specific probes such as dimethyl sulfate (DMS), 1-cyclohexyl-(2-morpholinoethyl)carbodiimide metho-p-toluene sulfonate (CMCT), and kethoxal [14], 2) backbone-cleaving reagents such as hydroxyl radicals [15] and metal ions [16], and 3) SHAPE (selective 2’-hydroxyl acylation analyzed by primer extension) reagents that modify the 2’-OH of the RNA backbone [12,17].

Chemical probes of RNA structure react with specific nucleotide positions that are solvent-accessible, or flexible, to covalently modify them. Modification positions are typically mapped with reverse transcription (RT), which either stops at these positions [12,14] or introduces a mutation into the cDNA [18]. An analysis of the resulting cDNAs can then be used to determine modification frequency at each nucleotide position. Modern approaches couple chemical probing and RT with next-generation sequencing (NGS) to directly sequence the cDNA products and determine modification positions [18-27]. The use of next generation sequencing mmediately enables these experiments to be highly multiplexed, allowing up to thousands of RNAs to be probed and analyzed in a single experiment. These extensive datasets [28] can be analyzed in many ways [11] and are routinely used to restrain RNA structure prediction algorithms [29].

The recent transition to NGS-based methods has also been accompanied by a second transition in the field: the move from characterizing RNA structure *in vitro* in favor of *in vivo* [11], Chemical probing *in vivo* requires that the probe be able to quickly diffuse across membranes to modify RNAs inside the cell. While DMS has long been known to penetrate cell membranes [30], it was also recently shown that some members of the SHAPE family of reagents can do so as well [31,32], Shortly after the first NGS-based chemical probing method was published 19], a number of different approaches were taken to combine NGS with *in vivo* or *in virio* chemical modification [18,21-23,26,27], many of which were to designed to probe the entire transcriptome of their respective organism [21-23,27].

The growing number of NGS-based techniques have many basic steps in common including chemical modification, reverse transcription, and PCR for library preparation (Figure 1) [18,20-23,26,27]. However, there are many details involved in several steps of these methods that present challenges. Beyond the continuous need for greater sequencing depth at lower costs, reducing the time, effort, complexity, and cost of the NGS library preparation steps are the biggest barriers to wide adoption of NGS-based chemical probing methods. For example, many NGS-based methods use gel purification steps that are typically time intensive and reduce cDNA yield. In previous work, we simplified the process for *in vitro* studies by describing SHAPE-Seq v2.0 [20], which reduced library preparation time and added a ‘universal’ RNA ligation method to characterize short RNA sequences by priming RT from a 3’ ligated linker sequence post-modification. We also developed in-cell SHAPE-Seq to characterize the structures of small groups of targeted RNAs directly inside cells [26]. Although not originally designed to cover the entire transcriptome, in-cell SHAPE-Seq avoids gel purification steps, making it much quicker and more amenable to first time users of *in vivo* NGS-based chemical probing methods. Overall, in-cell SHAPE-Seq is an approachable technique for RNA biologists interested in studying a few select RNAs rather than the whole transcriptome [26].

**Figure 1.**
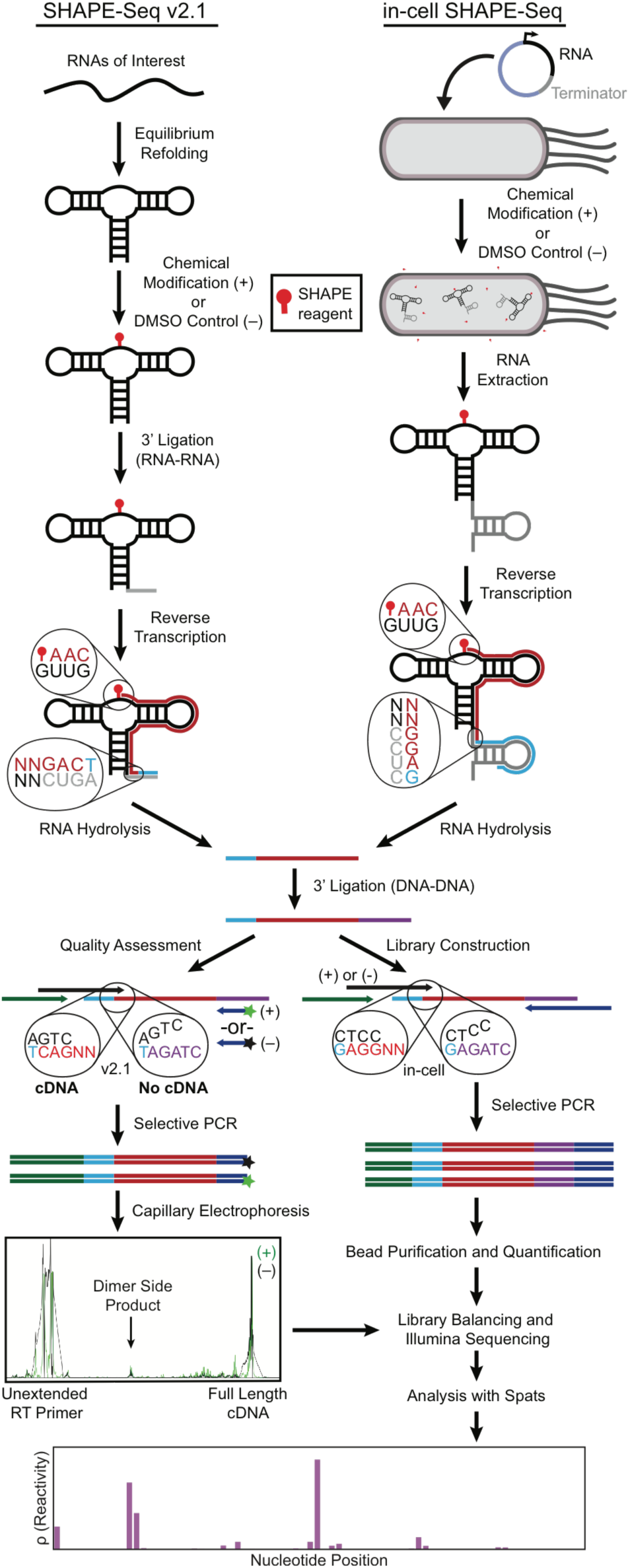
SHAPE-Seq workflow. *in vitro* RNA structures are analyzed by first purifying RNAs of interest, refolding in an appropriate buffer with optional ligands, and modifying with a SHAPE reagent (+) or a control solvent (-). In-cell probing experiments modify RNAs within the cell after the SHAPE reagent or control solvent is added to the media. Reverse transcription (RT) is initiated in one of two ways. In SHAPE-Seq v2.1, an RNA linker sequence is ligated to RNAs that serves as an RT priming site, whereas in-cell probed RNAs are extracted then primed directly with an internal RT priming site, such as the intrinsic terminator shown. Insets show specific 5’ and 3’ cDNA sequences, the latter of which is used to identify the SHAPE modification position. After reverse transcription, all SHAPE-Seq steps are similar. After RNA hydrolysis, a DNA adapter required for Illumina sequencing is ligated to the 3’ cDNA ends. Selective PCR is then used to generate quality assessment or sequencing libraries. The selective PCR uses a selection primer (black) designed to bridge the RT priming site and the 5’ end of the extended cDNA. This allows efficient PCR amplification only if the RT reaction created cDNA extensions. If no cDNA was synthesized, the selection primer cannot bind properly to the RT primer-adapter side product junction (inset), limiting amplification. Quality assessment libraries use a reverse primer that is fluorescently labeled for analysis with capillary electrophoresis, while sequencing libraries use the full Illumina adapter sequence. Subsequent sequencing and bioinformatic analysis (Spats) of the (+) and (-) libraries generates the characteristic SHAPE-Seq reactivity spectra.

In this work, we outline approaches for characterizing RNA structure *in vitro* and *in vivo* using SHAPE-Seq v2.1 and in-cell SHAPE-Seq, respectively. SHAPE-Seq v2.1 further upgrades the v2.0 technique to incorporate more flexible library barcoding capabilities and a re-engineered linker sequence to permit selective amplification of cDNA sequences as inspired by in-cell SHAPE-Seq [26]. In addition to describing experimental aspects of these techniques, we cover computational approaches that incorporate the resulting SHAPE-Seq reactivity data to improve the prediction of RNA structure models. We also highlight how comparing reactivity differences with and without ligand or *in vitro* vs. *in vivo* can reveal information about the effects of the folding environment on RNA structure. We discuss important considerations for performing SHAPE-Seq experiments and restraining computational predictions. Finally, we also suggest future improvements for the SHAPE-Seq technique and the interpretation of its reactivity data.

## 2. SHAPE-Seq Background

The SHAPE-Seq protocol consists of several core steps. These include: RNA chemical modification, converting RNA into cDNA with reverse transcription, sequencing the cDNA, bioinformatically processing sequencing reads, and, optionally, using the reactivities to restrain computational folding algorithms (Figure 1). Here we discuss the approaches and relevant background for each step before covering them in greater detail in Section 3.

### 2.1 RNA modification

To begin, the RNA of interest and its proper folding environment need to be determined. For *in vitro* studies, this involves determining the proper folding buffer and conditions (times, temperatures, ligand concentrations, etc.). For *in vivo* studies, the main choices to consider are which organism is being examined and whether endogenous or exogenous RNAs (e.g. plasmids, etc.) are being targeted. Once determined, the RNA is first folded in the chosen environment then treated with a SHAPE reagent (+), or a solvent control (-), by adding it directly to the *in vitro* solution or the cell culture media (Figure 1). The modified RNAs are then prepared for reverse transcription (RT) according to the type of experiment being performed. For example, *in vivo* studies, or *in vitro* studies with proteins, first require a two-phase extraction to remove proteins and/or contaminating DNA. For *in vitro* experiments where internal RT priming is not convenient, a ligation step introduces extra RNA sequence at the 3’ end to serve as an RT priming site [20]. In all cases, the final RNA form is concentrated using ethanol precipitation before RT.

### 2.2 Conversion of RNA to cDNA with reverse transcription

For both *in vitro* and *in vivo* experiments, reverse transcription and the steps that follow are largely the same (Figure 1). An RT primer specific to either an internal RNA sequence or ligated linker is added for extension with reverse transcriptase, which stops one nucleotide before the SHAPE modification position. The RNA is then hydrolyzed, followed by ethanol precipitation to wash away the base. Next, a DNA adapter is added to the 3’ end of the cDNA via a single stranded DNA-DNA ligation. The DNA adapter introduces one of the Illumina sequencing adapters required for DNA amplification and downstream sequencing.

This DNA-DNA ligation step is fairly standard among NGS-based chemical probing techniques [11,19-27]. It is also one reason that many NGS-based methods require gel purification, as the ligation products tend to be a complex mixture of oligonucleotides that contains a large amount of unwanted side product or starting material that needs to be removed before sequencing library preparation. One recently described solution to the purification problem has been to include an azide group on the modifying reagent, which can then be covalently linked to a biotin moiety via a ‘click’ reaction for selective pull-down [27].

Alternatively, as part of our in-cell SHAPE-Seq technique, we developed a selective PCR step that only allows significant amplification of correctly ligated products containing some length of transcribed cDNA (Figure 1) [26]. Selective PCR removes the need for gel purification and reduces the time, cost, and expertise required to prepare sequencing libraries. As described below, we also altered the SHAPE-Seq v2.0 linker to allow for selective PCR, which was not part of the original v2.0 protocol [20]. Finally, in all SHAPE-Seq methods a final bead purification step is employed to reduce the amount of unligated adapter present.

### 2.3 Preparation for sequencing

To prepare libraries for sequencing, the ssDNA libraries from Step 2.2 are amplified with PCR to add the complete Illumina TruSeq adapter sequences on each end that are required for sequencing. The adapters contain DNA sequences necessary for binding to the flow cell, priming the sequencing reactions, and barcoding the SHAPE-Seq libraries to sequence multiple libraries on a single flow cell. The PCR step requires three oligonucleotides (Figure 1). The first is the reverse primer that binds to the ligated DNA adapter and adds a TruSeq index and one of the flow cell binding sequences. The second primer contains the other flow cell binding sequence and the sequencing primer site for the first read of sequencing (RD1). The third, or selection primer, selects against unwanted DNA adapter-RT primer ligation side products during PCR and consists of a combination of the RD1 sequence and a designed sequence specific to the RT primer that extends 2-5 nt into the cDNA (Figure 1). In this PCR amplification scheme, the ligation side product formed between the unextended RT primer and the DNA adapter cannot be exponentially amplified due to a 3’ overhanging mismatch, providing a mechanism of selection against this side product, which can be present at a high concentration.

After PCR library construction, DNA library quality can be determined using either an Agilent BioAnalyzer or similar equipment to verify that the length distribution of the library matches the expected lengths for the RNA(s) tested. Alternatively, we prefer a more sensitive quality control reaction, where the reverse primer is replaced with a shorter, fluorescently labeled version for visualization with capillary electrophoresis (Figure 1).

Once the library quality is verified, individual libraries containing different TruSeq indexes are measured for concentration and pooled before sequencing with either an Illumina MiSeq or HiSeq instrument, using short paired-end reads.

### 2.4 Bioinformatic read alignment and reactivity calculation

Sequencing data is converted to chemical reactivity values by first aligning all of the sequences to the RNA(s) being studied to establish profiles of stop frequency for the modified RNA sample (+) and the unmodified control (-). These profiles are then used in a maximum likelihood estimation procedure to determine the relative modification frequency, or reactivity, of each nucleotide in the RNAs [33,34] (see Section 5.6). High reactivities suggest unpaired nucleotides, while low reactivities are indicative of structural inflexibility due to base pairing, helical stacking, tertiary contacts, protein interactions, or other factors that can reduce nucleotide flexibility [35-37].

### 2.5 RNA structure prediction using SHAPE-Seq reactivities

A major application of RNA chemical probing is to use reactivity data as restraints in computational RNA folding algorithms to improve structural predictions *in silico*. In these methods, users supply an RNA sequence along with reactivity data as inputs to generate predicted RNA structures that are more consistent with the experimental data. Several such methods incorporating reactivity values have shown that the use of SHAPE reactivities improves RNA structure prediction accuracy [20,38-41].

There are two main approaches for restraining computational RNA folding algorithms with SHAPE-Seq reactivities: 1) directly modifying the folding calculation or 2) selecting the structure from the results of the folding calculation that is most consistent with the experimental data. Both methods first calculate a partition function that describes how a population of RNA molecules partitions into an ensemble of different structures in equilibrium, with each structure occurring with a distinct probability [42]. Many properties can be determined from the partition function, including the minimum free energy (MFE) structure, which has the highest probability of occurring in the ensemble.

The first method to directly modify the RNA folding calculation was introduced by Deigan *et al*. as part of the RNAStructure suite of tools [38]. To use experimental SHAPE reactivities in the folding calculation, they are first converted into pseudo-free energy terms Δ*G_SHAPE_* (*i*) that are included for each nucleotide *i* in the calculation of the RNA structure’s overall free energy (Δ*G*) according to:

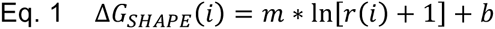

where *r*(*i*) is the reactivity at nucleotide *i*, and *m* and *b* are parameters that were originally fit by comparing a restrained prediction to the known crystal structure of the 16S rRNA from *E. coli* [38]. By including Δ*G_SHAPE_*(*i*) in the free energy calculation when the nucleotide *i* is paired, MFE structures are returned that are more consistent with the observed reactivity data [20,38,43]. Work has also been done to predict pseudoknot interactions using *ShapeKnots*, an algorithm that runs two folding stages that include Δ*G_SHAPE_*(*i*) and another pseudo-free energy term for pseudoknots [39].

The second method uses the RNA structure partition function to generate a set of possible RNA structures [44] and then selects the structure(s) that most closely agree(s) with the experimental data. To perform this type of selection, the algorithm SeqFold first converts SHAPE reactivities to a “structural preference profile” (SPP), a normalized vector of reactivities restricted to [0,1] [45]. Each RNA structure is similarly converted into a binary vector such that 0 and 1 represent paired or unpaired bases, respectively. A minimum distance structure is then chosen by calculating the distance between the SHAPE SPP and each possible structure vector. Finally, the cluster of structures most closely related to this minimum distance structure is used to calculate a representative centroid structure [45]. Unlike MFE-based methods, this cluster-based approach provides more information about different structure sub-populations.

Recently, a method that combines both approaches was developed called restrained MaxExpect (RME). RME uses SHAPE reactivities to modify the partition function and selects a structure from it that best matches the experimental data [46]. First, RME calculates a partition function after adding a pseudo-free energy term for each nucleotide *i* using:

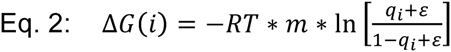

where *m* is the weighting parameter for the pseudo-free energies, *ε* is a small constant value to ensure a real answer, and *q_i_* is the measured base pair probability obtained from the experimental reactivities for position *i*, assigned to a ‘low’ or ‘high’ value of 0 or 1 based on a reactivity cutoff to represent paired or unpaired bases, respectively. Then, a predicted base pairing probability matrix is derived from the partition function that is then adjusted by the experimental data to account for discrepancies between the predicted base pairing matrix and the measured reactivities. This newly modified base pairing matrix is finally used to predict a structure that maximizes expected accuracy [46].

This concept of modifying the results of a partition function calculation with experimental data was introduced previously by Washietl *et al*. [47]. In their method, the free energy calculation for each RNA structure is perturbed with pseudo-free energy terms that are numerically determined instead of explicitly calculated (e.g. Eq. 1). The method first estimates an experimental base pairing vector using a reactivity cutoff as in RME. Then a partition function calculation is performed in which the free energy model is perturbed with a vector of pseudo-free energy terms for each nucleotide in the RNA. This vector of pseudo-free energies is adjusted iteratively to minimize the differences between the predicted base pairing probability matrix and the estimated experimental base pairings [47]. After arriving at the final base pair probability matrix, it is used to predict a structure that maximizes expected accuracy as with the RME method above.

Finally, Kutchko *et al*. took a different approach to RNA structure prediction by examining ensembles of structures through their subpopulations rather than picking or predicting a single structure for each set of SHAPE reactivities [48]. In this method, SHAPE reactivities are used to modify the partition function calculated by RNAStructure using the free energies introduced in Eq. 1. The modified partition function is then used to generate many possible structures that are converted into binary vectors for structure clustering as described above. The resulting structures near cluster centroids represent different RNA structure subpopulations in the ensemble that are more consistent with the experimental data [48]. The benefit of this approach is that the entire ensemble of structures is analyzed rather than focusing on a single ‘best’ predicted structure.

## 3. Materials and Methods

### 3.1 RNA folding and modification

The following steps outline RNA folding and modification with 1-methyl-7-nitroisatoic anhydride (1M7) [49], which can be synthesized in a one-step reaction [50]. Alternatively, N-methylisatoic anhydride (NMIA) can be commercially purchased and used in these steps with the noted modifications.

#### 3.1.1 in vitro *experiments*

1. Generate dsDNA templates with the promoter sequence for T7 RNA polymerase followed by the DNA sequence encoding the RNA of interest for run-off transcription. For the v2.1 method, where an RT priming site will be added later with ligation (Step 3.2 below), we recommend following the RNA of interest with the hepatitis δ ribozyme to produce the correct 3’ end after cleavage and 3’ end healing [20,51]. Otherwise, the tendency of T7 RNA polymerase to add 1-3 spurious nucleotides to the 3’ end of the RNA can cause alignment issues downstream if not accounted for.
2. Set up an in vitro transcription reaction using standard methods [19,20,52].
3. Ethanol precipitate the transcription products to concentrate them.
4. Gel purify the RNA of interest. If UV shadowing is used, be careful to avoid directly shadowing the RNA being purified to avoid damaging the RNA [53].
5. Check RNA integrity using a polyacrylamide gel.
6. Choose an appropriate folding buffer for the RNA of interest and prepare a 3.3X concentrated solution. A good starting point folding buffer (1X) is: 10 mM MgCl_2_, 100 mM NaCl and 100 mM HEPES (pH 8.0) [52].
7. Dilute 1-20 pmol of RNA in 12 μL RNase-free H_2_O. Denature at 95 °C for 2 min. Snap cool on ice for 1 min, then add 6 μL of 3.3X buffer. Incubate at 37 °C for 20 min. If adding an RNA-binding protein, do so after the second incubation, and follow with a third incubation to give the protein time to bind. This step can be adjusted based on the folding conditions desired.
8. Prepare a 65 mM solution of 1M7 in anhydrous DMSO. Aliquot 1 μL each of 65 mM 1M7 and anhydrous DMSO into different tubes. Upon completing the RNA folding incubation, add 9 μL to the 1M7-containing tube (+ sample) and mix. Do the same for the other 9 μL with DMSO (- sample). Incubate for 2 min at 37 °C to complete the reaction [20]. See Section 5.3 below for modifications if other folding conditions or chemical probes are used.

#### 3.1.2 in vivo *experiments*

1. If the RNA of interest is not endogenously expressed, clone it into an expression vector, being sure to include a priming site for reverse transcription. Two examples of convenient RNA expression vectors containing specific RT primers for superfolder green fluorescent protein mRNA and a synthetic intrinsic terminator can be found in Watters *et al*. [26]. These vectors are available on Addgene.
2. Grow 1 mL of cells into the desired growth phase, increasing the growth volume if more culture will be required for a functional assay. For *E. coli* grown at 37 °C, an OD value within 0.3-0.8 is recommended for exponential phase. Induce RNA expression if required and allow an appropriate amount of time for RNA synthesis.
3. Prepare a 250 mM solution of 1M7 in anhydrous DMSO. Aliquot 13.3 μL each of 250 mM 1M7 and DMSO into different tubes. See Section 5.3 below for modifications to this step if other chemical probes are used.
4. If taking a functional measurement (e.g. a fluorescence assay to determine regulatory function of a non-coding RNA [26]), set aside the volume of cell culture required for the assay, leaving 1 mL for RNA modification. For *E. coli*, we suggest pelleting the functional test aliquot (typically 150 μL) and resuspending in cold PBS with antibiotics to prevent further gene expression [26]. At the same time, add 500 μL of cell culture into each tube of 1M7 or DMSO and mix. Incubate at the culture growth conditions with shaking for 2-3 min to complete the reaction.
5. Extract the total RNA quickly to prevent significant RNA degradation. Any RNA isolation method is acceptable. For *E. coli*, we recommend the TRIzol Max Bacterial RNA Isolation Kit (ThermoFisher) [26]. Dissolve/Elute the extracted total RNA with 10 μL of RNase-free H_2_O.

### 3.2 *RNA linker ligation (skip for* in vivo *or direct priming experiments)*

1. Prepare 5’-adenylated linker by purchasing the phosphorylated RNA linker sequence (5’App-CUGACUCGGGCACCAAGGA-3’ddC) from an oligonucleotide supplier and the 5’ DNA Adenylation Kit from New England BioLabs (NEB). Follow the manufacturer’s instructions to adenylate 500 pmol of RNA linker in 50–100 uL reaction aliquots. Purify the RNA linker with a phenol-chloroform or TRIzol (ThermoFisher) extraction. Quantify the amount of RNA after purification and prepare a 20 μM solution of the purified 5’-adenylated RNA linker.
2. Ethanol precipitate the modified RNA from the end of Step 8 of Section 3.1.1. If proteins and/or DNA were present in the folding conditions, perform a phenol-chloroform or TRIzol extraction first. Dissolve the pellet in 10 μL of 10% DMSO in RNase-free H_2_O.
3. Mix the ligation reaction by adding: 0.5 μL of SuperaseIN (ThermoFisher), 6 μL 50% PEG 8000, 2 μL 10x T4 RNA Ligase Buffer (NEB), 1 μL of 20 μM 5’-adenylated RNA linker, and 0.5 μL T4 RNA Ligase, truncated KQ (NEB). Lower concentrations of the adenylated linker can be used as long as the linker is in at least 4-fold excess. Mix well and incubate overnight at room temperature.
4. Ethanol precipitate the RNA using glycogen as a carrier and dissolve the pellet in 10 μL of RNase-free H_2_O.

### 3.3 Reverse transcription

1. Add 3 μL of 0.5 μM RT primer to the dissolved RNA. For *in vitro* experiments with ligation, the RT primer sequence should be GTCCTTGGTGCCCGAGT. For internally primed reactions, an appropriate RT primer should be designed prior to this step and used for this reaction.
2. Prepare the reverse transcription master mix by combining 0.5 μL of Superscript III (ThermoFisher), 4 μL 5X First Strand Buffer (ThermoFisher), 1 μL 100 mM dithiothreitol (DTT), 1 μL 10 mM dNTPs, and 0.5 μL RNase-free H_2_O.
3. Incubate the RNA and RT primer at 95 °C for 2 min, followed by 65 °C for 5 min. Snap-cool the tubes for 30 seconds, then add 7 μL of the master mix from Step 2 and mix well.
4. Incubate at 52 °C for 25 min, followed by 65 °C for 5 min to inactivate the reverse transcription. These RT conditions may be adjusted if necessary for specific RNAs that are difficult to reverse transcribe.
5. Hydrolyze the RNA by adding 1 μL of 4 M NaOH to the RT reactions and incubate at 95 °C for 5 min (Replace 4 M NaOH with 10 M NaOH for *in vivo* experiments). Add either 2 μL or 5 μL of 1 M HCl for *in vitro* or *in vivo* experiments, respectively, to partially neutralize remaining base.
6. Ethanol precipitate the cDNA, washing the pellet thoroughly with 70% ethanol. Dissolve the pellet in 22.5 μL of nuclease-free H_2_O.

### 3.4 Sequencing adapter ligation

1. To the cDNA, add 3 μL 10X CircLigase Buffer (Epicentre), 1.5 μL 2.5 mM MnCl_2_, 1.5 μL 50 μM ATP, 0.5 μL 100 μM DNA adapter (5’-Phos-AGATCGGAAGAGCACACGTCTGAACTCCAGTCAC-3C Spacer-3’), and 1 μL CircLigase I (Epicentre). Mix well. (Note: the DNA adapter should be PAGE purified before use).
2. Incubate at 60 °C for 2 hr, then 80 °C for 10 min to inactivate the ligase.
3. Ethanol precipitate the ligated cDNA using glycogen as a carrier and dissolve in 20 μL of nuclease-free H_2_O.
4. Purify using 36 μL of Agencourt XP Beads (Beckman Coulter), according to manufacturer’s instructions to remove excess DNA adapter. Elute from beads with 20 μL TE buffer. The ssDNA libraries can now be stored until sequencing or quality analysis is performed.

### 3.5 Quality analysis

1. Design a pair of selection PCR primers that correspond to each RT primer used. To design the primer pairs, start with either the sequence CTTTCCCTACACGACGCTCTTCCGATCTYYYR (-samples) or CTTTCCCTACACGACGCTCTTCCGATCTRRRY (+ samples) as the 5’ end of the selection primer sequence. Then, append the desired RT sequence to the 3’ end of these sequences. Next, extend the RT primer sequence at the 3’ end to contain a few bases of the cDNA and protect these bases with phosphorothioate modifications to prevent 3’->5’ exonuclease degradation. As an example, the selection primer pair for samples that use the v2.1 *in vitro* linker ligation strategy are CTTTCCCTACACGACGCTCTTCCGATCT(RRRY/YYYR)GTCCTTGGTGCCCGAG*T*C *A*G, where * represents a phosphorothioate modification.
2. Mix a separate PCR reaction for each (+) and (-) sample by combining: 13.75 μL nuclease-free H_2_O, 5 μL 5X Phusion Buffer (NEB), 0.5 μL 10 mM dNTPs, 1.5 μL of 1 μM labeling primer (5’-Fluor-GTGACTGGAGTTCAGACGTGTGCTC-3’; see below), 1.5 μL of 1 μM primer PE_F (AATGATACGGCGACCACCGAGATCTACACTCTTTCCCTACACGACGCTCTTCCGATCT), 1 μL of 0.1 μM selection primer (+ *or* -) from Step 1, 1.5 μL ssDNA library (+ *or* -), and 0.25 μL Phusion DNA polymerase (NEB). Use two different compatible fluorophores for the (+) and (-) samples. We recommend VIC and NED, respectively. See Figure 1 for a schematic of this step.
3. Amplify PCRs for 15 cycles using an annealing temperature of 65 °C and an extension time of 15 seconds. More cycles can be used if the input RNA was low. However, when using a large amount of cycles, we recommend excluding primer PE_F from the reaction until the last 10–15 cycles to reduce side product formation [26].
4. Combine the (+) and (-) reactions, add 50 nuclease-free H_2_O, and ethanol precipitate. Dissolve the combined reactions in formamide and examine with capillary electrophoresis, looking for good full-length RT extension and low dimer side product (Figure 1). See Watters *et al*. for more details [26].
5. (Alternate method) Skip Steps 2–4 above and follow the steps for sequencing library preparation in Section 3.6 below. Check libraries on an Agilent BioAnalyzer or similar instrument, looking for good full-length RT extension and low dimer side product.

### 3.6 Library preparation for sequencing

1. Assess whether the libraries are of sufficient quality to sequence.
2. Mix a separate PCR for each (+) and (-) sample by combining: 33.5 μL nuclease-free H_2_O, 10 μL 5X Phusion Buffer (NEB), 0.5 μL 10 mM dNTPs, 0.25 μL of 100 μM TruSeq indexing primer (CAAGCAGAAGACGGCATACGAGATxxxxxxGTGACTGGAGTTCAGACGTGTGCTC; see below), 0.25 μL of 100 μM primer PE_F (Section 3.5 Step 2), 2 μL of 0.1 μM selection primer (+ *or*-, Section 3.5 Step 1), 3 μL ssDNA library (+ *or* -), and 0.5 μL Phusion DNA polymerase (NEB). Replace the ‘xxxxxx’ sequence with the appropriate six nucleotide TruSeq indexing barcode for Illumina sequencing. Additional barcoding can be added 5’ of the RT primer sequence in the selection primer pair, although these barcodes will not be detected and split automatically by the Illumina sequencing processing pipeline. See Loughrey *et al*. for more details [20].
3. Amplify PCRs for 15 cycles using an annealing temperature of 65 °C and an extension time of 15 seconds. If a different number of cycles was used during quality analysis, use that PCR configuration instead.
4. Add 0.25 μL of Exonuclease I and incubate at 37 °C for 30 minutes to remove excess primer. Allow the PCRs from Step 3 to cool before adding the enzyme.
5. Purify reactions using 90 μL of Agencourt XP Beads (Beckman Coulter), according to manufacturer’s instructions. Elute from the beads with 20 μL TE buffer. The dsDNA libraries are complete and ready for sequencing.

### 3.7 Illumina sequencing

1. Determine the mass concentration of all dsDNA libraries to be sequenced. We recommend using the Qubit high sensitivity DNA kit (ThermoFisher). Then calculate the molarity after determining the average molecular weight using the average length of the dsDNA library from the quality analysis traces [26].
2. Choose either the MiSeq or HiSeq platforms for sequencing. As a conservative rule of thumb, we suggest running one library pair (+ and -) for each million reads provided by the sequencing kit chosen. However, more libraries can be sequenced at once, provided that the amount of unwanted dimer side product is low.
3. Sequence the library mixture according to the manufacturer’s instructions using 2x35 bp paired-end reads. Significantly longer read lengths are not recommended or necessary.

### 3.8 Data analysis with Spats

1. Obtain the compressed sequencing data on a Linux, Unix, or Mac OS X capable computer and extract the “.fastq.gz” files. Each TruSeq index will contain a pair of sequencing data files generated from the Illumina sequencing processing pipeline.
2. Create a fasta (.fa) formatted targets file that contains all of the RNA sequences that were measured in each TruSeq index. Include all of the linker sequences, internal barcodes, etc. if present.
3. Download and install the latest version of Spats (https://github.com/LucksLab/spats), including its utility scripts and dependent programs. Detailed instructions for running Spats and its utility scripts can be found in Watters *et al*. [26].
4. Run adapter_trimmer.py with the following command: *adapter_trimmer.py <R1_seq.fastq> <R2_seq.fastq> <targets.fa>* where <R1_seq.fastq> and <R2_seq.fastq> are the Illumina “.fastq” files for Read 1 (R1) and Read 2 (R2), respectively, and <targets.fa> is the fasta-formatted targets file created in Step 2.
5. Run Spats on the output file pair from adapter_trimmer.py using the following command: *spats <targets.fa> RRRY YYYR combined_R1.fastq combined_R2.fastq*
6. Normalize the output θ_i_ values to ρ_i_ values by multiplying all of the θ_i_ values by one less than the original RNA length [20,26] (See Section 5.6 below). Do not include the linker or adapter sequences in this length. The ρ reactivities can be used to restrain secondary structure folding (See Section 3.9 below).

### 3.9 SHAPE-directed computational RNA folding

In the following steps we outline the process for restraining three different RNA folding algorithms with SHAPE-Seq reactivity data: RNAStructure [39,54], restrained MaxExpect (RME) [46], and the Washietl *et al*. method (as part of the RNAprobing webserver) [47]. As discussed in Section 2.5, RNAStructure (containing Fold and ShapeKnots) can calculate the MFE structure directly as well as generate an ensemble of structures (with the *partition* and *stochastic* commands).

#### 3.9.1 Algorithm 1: RNAStructure

1. Install the RNAStructure text interface program (http://rna.urmc.rochester.edu/rnastructure.html) on a Linux, Unix, or Mac OS X operating system [54]. Alternatively, the webserver tools are available at http://rna.urmc.rochester.edu/RNAstructureWeb/.
2. Create a sequence file with a “.seq” extension that contains the following individual lines in order: 1) RNA name, 2) RNA sequence followed by the number ‘1’ to indicate the end of the file. Comment lines may be added as well, if preceded by a semicolon at the beginning of the file. Note that within the RNA sequence lowercase type forces the base to be single-stranded in the folding calculation. Uppercase denotes bases to include in the folding calculation. The letter ‘T’ is used as ‘U’ for RNA predictions.
3. Create a SHAPE reactivities file with a “.shape” extension that contains two tab-separated columns. The first column is the nucleotide number, starting with one, and the second column is the reactivity value for that position calculated by spats (ρ).
4. To run the *Fold* algorithm [55] (pseudoknots forbidden) use the command: *Fold <.seq file> <.ct file> -sh <SHAPE file> -m 1 -sm 1.1 -si “-0.3”* To run *ShapeKnots* [39] (up to one pseudoknot allowed) use the command: *ShapeKnots <.seq file> <.ct file> -sh <SHAPE file> -m 1 -sm 1.1 -si “-0.3”* For both commands, replace <.seq file> and <SHAPE file> with the files created in steps 2 and 3, respectively. In order, the options ‘m’, ‘sm’, and ‘si’ options correspond to the number of structures drawn, the SHAPE slope parameter *(m* in Eq. 1), and the SHAPE intercept parameter *(b* in Eq. 1). The values of 1.1 for *m* and -0.3 for *b* were fit for SHAPE-Seq *ρ* inputs in Loughrey *et al*. [20]. The output is *<.ct file>*, which needs to be specified as a “.ct” filename in the command. Once generated by the algorithm, it contains the minimum free energy structure as predicted by *Fold* or *ShapeKnots*. The first line of the “.ct” file will contain the length of the sequence, the free energy of the structure, and the title of the structure, respectively. The following lines contain, from left to right: the number of nucleotide *i*, its base, *i* - 1, *i* + 1, the number of its base pair partner (0 if unpaired), and the natural numbering (typically t).
5. As an alternative to *Fold* or *ShapeKnots*, RNAStructure can also sample structures from a partition function using the commands *partition* and *stochastic* [48]. Calculate the partition function based on SHAPE reactivities with the command: *partition <.seq file> <.pfs file> -sh <.shape file> -sm 1.1 -si “-0.3”* All of the options are the same as in Step 4 above, except <.pfs>, which is the calculated partition function output file. This partition function can then be used to sample structures using: *stochastic <.pfs file> <.ct file> -e <#> -seed <random integer>* The above command will sample the number of structures specified by the *-e* option and output them in a concatenated list to the *<.ct file>*. Changing the *-seed* option (default of 1234) will result in a different set of sampled structures.

#### 3.9.2 Algorithm 2: RME

1. Install R version 3.2.2 (https://www.r-project.org/) and the packages rshape, mixtools, and evir. Also install Bioconductor (http://bioconductor.org/install/) and the RME source code, found at https://github.com/lulab/RME [46].
2. Create a FASTA file of the RNA sequence to analyze.
3. Create a “.ct” file for the RNA sequence being analyzed using *Fold* (Step 4 in Section 3.9.1). Note that the base pairing information is not used during the RME calculation.
4. Prepare a tab-separated data file containing the reactivity information. The first line should read “RNA<tab>Index<tab>Reactivity<tab>Base”. The following lines should contain the RNA name, index, reactivity, and base (e.g. TPP, 1, ρ_1_, G; TPP, 2, ρ_2_, C, etc.) for the entire length of the RNA.
5. Locate the SHAPE example training and test files in the /example/dat/data/ directory. They are required for calculation. Copy the “.ct” files from /example/dat/structure/ to the directory containing the “.ct” files generated in Step 3.
6. Pre-process the SHAPE data using the 23S rRNA training data according to: *RME-Preprocess -d SHAPE -s <ct files directory>* *example/dat/data/SHAPE.train.23SrRNA. data example/dat/data/SHAPE. test. data <pre-process directory>* where <pre-process directory> is a specified directory name to store the pre-processing data.
7. Predict SHAPE-directed structures using the following commands: *RME -d SHAPE <pre-process directory>/SHAPE.for-test.txt SHAPE* where <prediction directory> is a supplied directory name for the folding output and <pre-process directory> is the same as Step 6. The output will be in the in the form of a “.ct” file, which can be used in the same manner as the folding results from Section 3.9.1.

#### 3.9.3 *Algorithm 3: RNAprobing webserver (Washietl* et al. *method)*

1. Go to the RNAprobing WebServer at http://rna.tbi.univie.ac.at/cgi-bin/RNAprobing.cgi.
2. Enter the RNA sequence either by copy-paste or uploading a fasta formatted file.
3. Upload a SHAPE reactivities file according to Step 3 of Section 3.9.1. Note that SHAPE reactivities of 0 must be input as “0” and not “0.0” in order to be parsed properly.
4. Select “Washietl et al. 2012” as the “SHAPE method” and “Cutoff” from the dropdown window for “Method used to derive pairing probabilities”. A cutoff threshold of 0.25 was used in Washietl *et al*. [47], although other values can be used.
5. Click “Proceed” and you’ll be redirected to an output page once the calculations are complete. The output will include the dot bracket notation of the predicted structure and an image of the predicted structure.

#### 3.9.4 Drawing secondary structures

The “.ct” results file generated from the final steps of Sections 3.9.1 and 3.9.2 can be converted to a “.dbn” file that contains the structure in dot bracket notation. One way to do this is using the RNAStructure command:

*ct2dot <ct file> 1 <bracket file>*

where <ct file> is the “.ct” file being converted and <bracket file> is the output file name ending in “.dbn”. Note that *ct2dot* removes pseudoknots that may be generated by *ShapeKnots*. A number of different programs, such as *VARNA* (http://varna.lri.fr/) [56] can use dot bracket information to draw the secondary structure of an RNA sequence. The *draw* command in RNAStructure [57] can also draw secondary structures using “.ct” files.

## 4. Results

### 4.1 in vitro *SHAPE-Seq analysis*

SHAPE-Seq v2.0 consisted of several protocol optimizations to simplify and shorten the original SHAPE-Seq technique [19,52]. In addition, it increased the technique’s flexibility by adding a 3’ linker ligation step after modification to remove RT priming site restrictions within the RNA [20]. In this work, we expand on these improvements by adapting the mismatch PCR selection developed in Watters *et al*. for in-cell SHAPE-Seq [26] to SHAPE-Seq v2.0 with a new redesigned linker for reduced dimer side product formation, thereby updating the *in vitro* experiment to SHAPE-Seq v2.1.

#### 4.1.1 SHAPE-Seq v2.0 vs. v2.1

We compared SHAPE-Seq v2.1 to v2.0 using two well-benchmarked RNAs [20,38,39,43]: 5S rRNA from *E. coli* and the *add* adenine riboswitch aptamer domain from *V. vulnificus*. For both RNAs, we folded and modified 40 pmol of RNA using the same buffer and ligand conditions as in Loughrey *et al*. [20] before splitting the (+) and (-) samples to process them individually with either the SHAPE-Seq v2.0 or v2.1 protocol. One immediately observable difference in the v2.1 vs. v2.0 raw sequencing data was a 25-fold reduction in the amount of DNA ligation side product sequenced with the v2.1 improvements. Additionally, we observed only slight differences between the reactivities obtained using v2.0 or v2.1, as expected (Figure 2). Further, the reactivity maps agree well with previous measurements [20,39,43].

**Figure 2.**
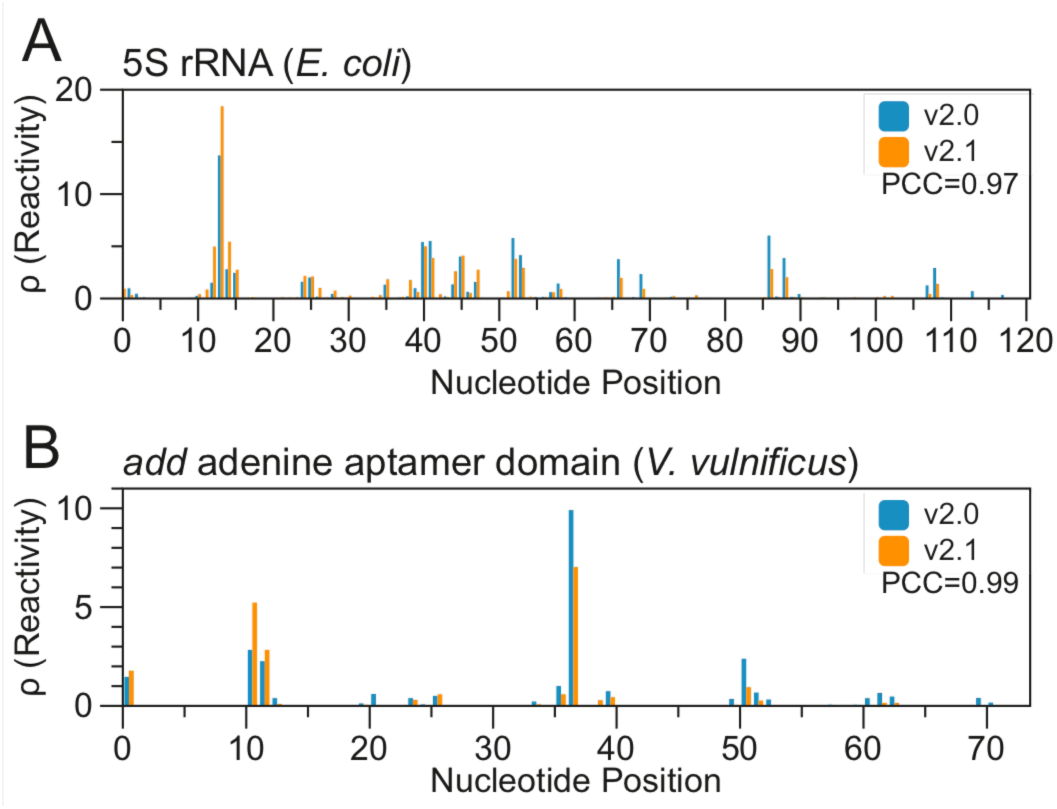
Comparison of SHAPE-Seq v2.0 vs. v2.1 *in vitro* reactivities. (A) Reactivity maps derived from the 5S rRNA from *E. coli* after equilibrium refolding and modification processed with either v2.0 (RMDB: 5SRRNA_1M7_0008) or v2.1 (RMDB: 5SRRNA_1M7_0009) library preparation steps. The reactivities closely agree, with a Pearson Correlation Coefficient (PCC) of 0.97. The ‘GGA’ sequence added to the 5’ end of the 5S rRNA to aid *in vitro* transcription is not shown, although included with PCC analysis. (B) The same analysis for the *add* adenine aptamer domain from *V. vulnificus* shows a PCC of 0.99 (RMDB: ADDsC_1M7_0007 and ADDSC_1M7_0008). Like the 5S rRNA, a ‘GG’ sequence was added to aid *in vitro* transcription and is not shown on the graph, but is included in the PCC analysis.

In general, reactivities derived from the v2.1 experiment tend to be slightly shifted toward the 5’ end of the RNA relative to v2.0 reactivities. This difference is likely due to the reduction of reads in the v2.1 data that align immediately upstream of the RT priming site in the area where the unwanted dimer side product appears. Thus, the reduction of these reads may represent a slightly more accurate reactivity map. However, this difference was minor, as we observed that the Pearson Correlation Coefficients (PCC) comparing v2.0 vs. v2.1 were in the range of 0.97–0.99 (Figure 2).

#### 4.1.2 Using SHAPE-Seq v2.1 to observe ligand binding

To demonstrate how SHAPE-Seq can be used to detect changes in RNA folding upon ligand binding, we examined the *thiM* thiamine pyrophosphate (TPP) riboswitch aptamer domain from E. *coli* using SHAPE-Seq v2.1. Comparing the reactivity maps with and without ligand for the TPP riboswitch aptamer domain shows a number of reactivity differences (Figure 3A).

**Figure 3.**
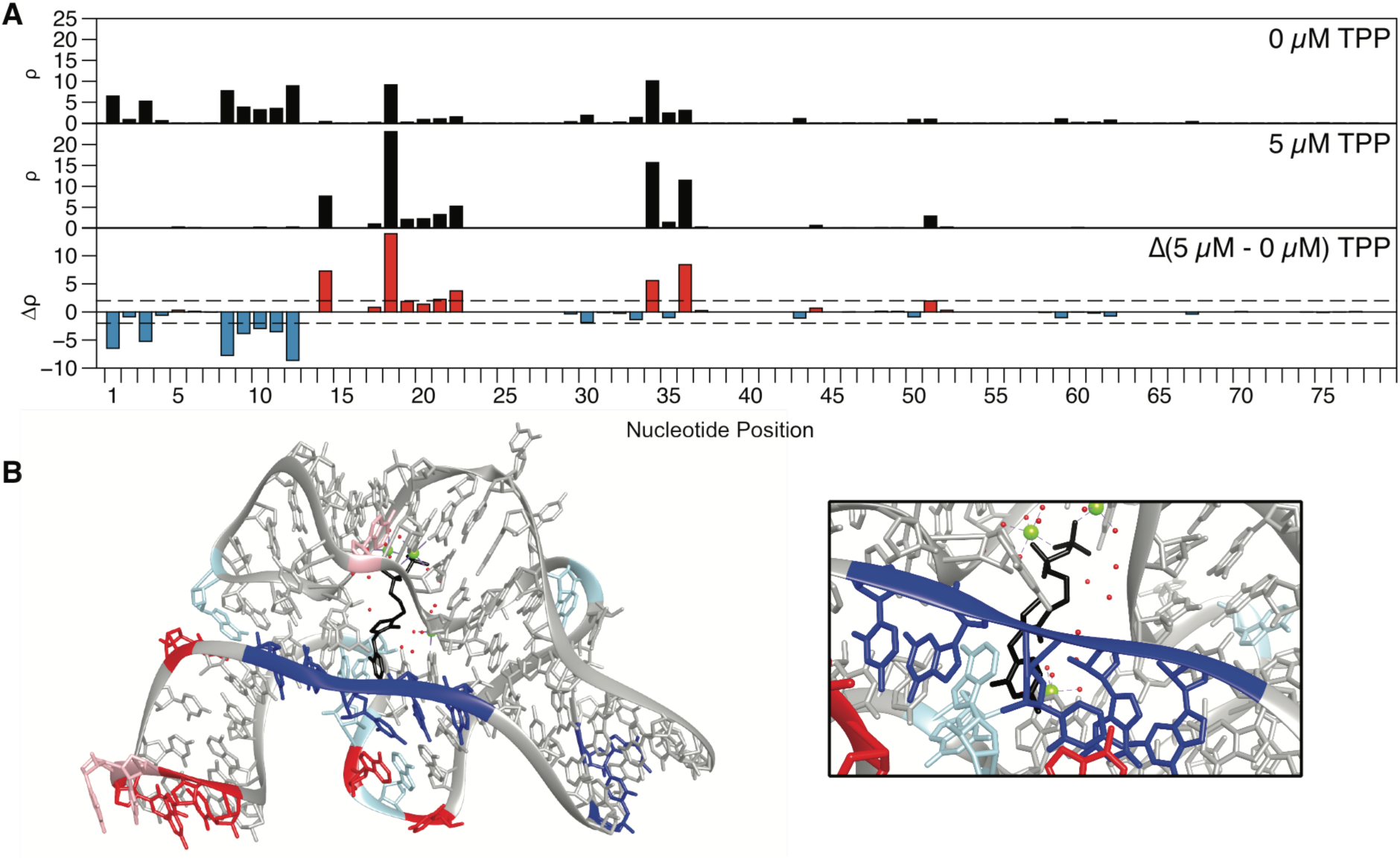
SHAPE-Seq reveals reactivity changes in the presence of ligand for the *thiM* TPP riboswitch aptamer domain. (A) *in vitro* reactivity maps for the *thiM* TPP aptamer domain with 0 μΜ or 5 μΜ TPP (RMDB: TPPSC_1M7_0005). The reactivity difference map (bottom) shows increases (red) and decreases (blue) in reactivity in the presence of ligand. (B) Crystal structure (pDB 2GDI) [60] of the *thiM* TPP aptamer domain with TPP (black) bound, colored by change in reactivity in the presence of ligand from part A. Magnesium ions are colored light green and the solvent is denoted as red dots. Nucleotides marked in red/blue show increases/decreases above |Δ*ρ*| ≥ 2 (dashed lines in part A). Light red and blue mark changes for which 1 < |Δ*ρ*| < 2. The region closing the TPP binding pocket shows a cluster of nucleotides that become less flexible upon ligand binding (inset). An extra 'G' was added to the 5' end to aid *in vitro* transcription and is not displayed.

Specifically, we observe an interesting set of decreasing reactivities at positions 8–12 near the binding pocket of TPP, suggesting that part of the ligand binding pocket is first flexible, but becomes rigid after ligand binding, which agrees with previous observations (Figure 3B; inset) [58]. These decreases also come with reactivity increases in positions 14, 17–22, 34, and 36.

#### 4.1.3 Inferring secondary structures with SHAPE-Seq data

We next investigated how incorporating SHAPE-Seq v2.1 reactivity values affects the calculated structures of the 5S rRNA, TPP riboswitch, and adenine riboswitch using the four methods described in Section 3.9. In general, we saw an improvement in both the percentage of known base pairs predicted correctly (sensitivity) and the percentage of predicted base pairs in the known structure (PPV; positive predictive value) for all of the methods used when SHAPE data was included (Figure 4), as has been shown multiple times [38,39,46,47,55].

**Figure 4.**
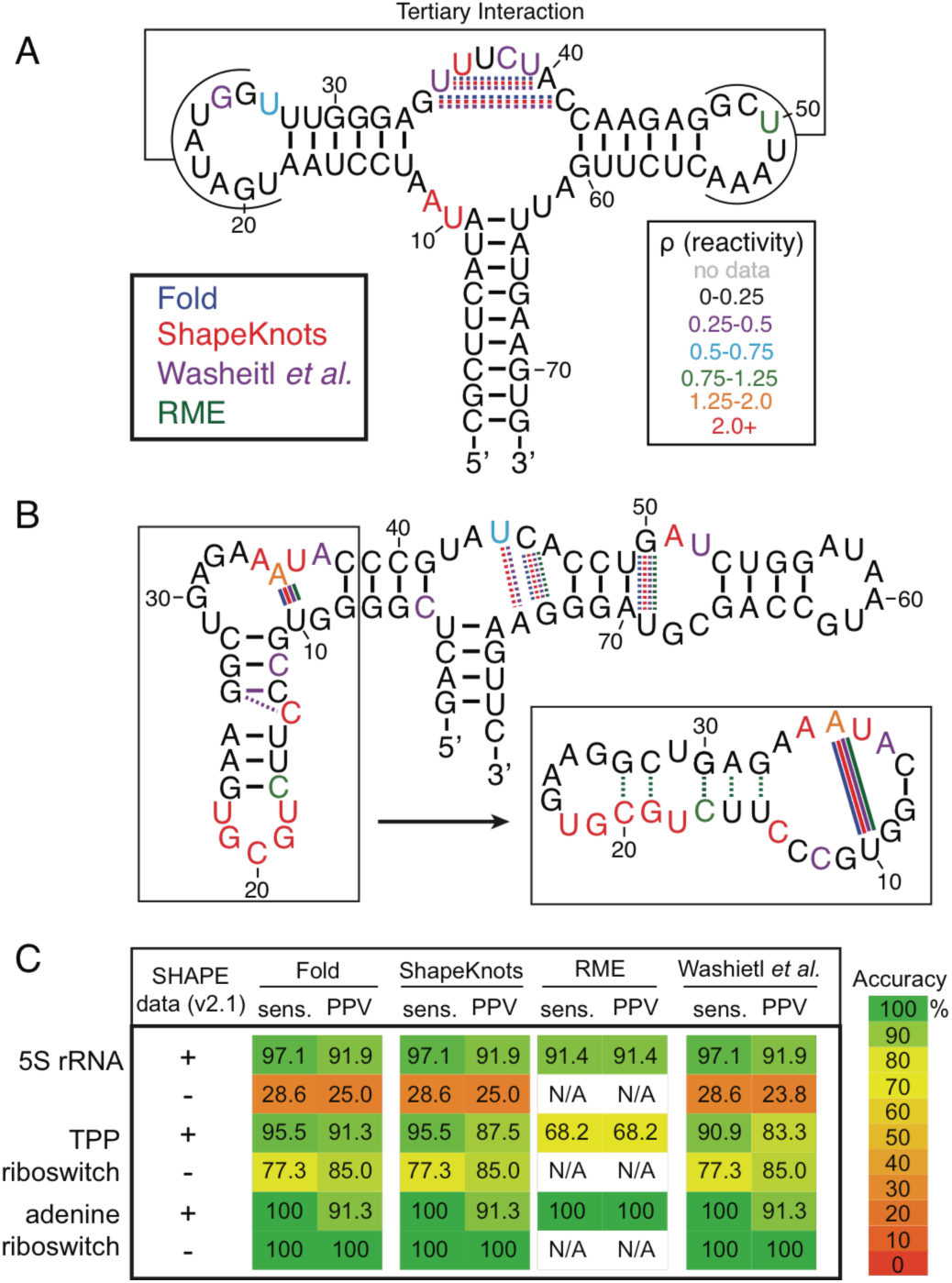
Incorporating SHAPE-Seq data improves computational folding accuracy. (A) *add* adenine riboswitch aptamer domain ligand-bound secondary structure representation from Serganov *et al*. [59]. Dashed lines mark base pairs predicted by computational methods, as indicated by color, restrained with SHAPE-Seq v2.1 reactivities (Figure 2B) that do not exist in the crystal structure representation. Colored solid lines indicate base pairs that are present in the crystal structure, but are not predicted with v2.1 reactivities. Individual nucleotides are color-coded by reactivity intensity. (B) *thiM* TPP riboswitch aptamer domain ligand-bound secondary structure representation from Serganov *et al*. [60]. Tertiary interactions and non-canonical base pairings are not shown. Solid and dashed lines represent the same features as in part A and reactivities (Figure 3A) are color-coded the same way. The predicted structure from RME is visibly different for nucleotides 8–38 (boxed) as drawn on the right. (C) Table summarizing the folding accuracies for the four computational algorithms *Fold* [38,55], *ShapeKnots* [39], Washietl *et al*. [47], and RME [46]. The Washietl *et al*. method was calculated using the RNAProbing webserver. No calculation could be performed for RME without SHAPE reactivities. Sens. = sensitivity, PPV = positive predictive value [42].

The adenine riboswitch folded into the expected ‘T’ structure with or without SHAPE data, although three of the methods predicted two extra base pairs relative to the secondary structure representation of the ligand-bound aptamer domain crystal structure (Figure 4A) [59]. The TPP riboswitch aptamer was also folded into the correct general structure in three of the four methods, although with some differences from the crystal structure representation (Figure 4B) [60]. It is worth noting, however, that for both RNAs most of the incorrect predictions occur in regions known to be involved in non-canonical base pairing or protein/ligand interactions that are represented as unpaired bases in the ‘accepted structure’ for the purpose of calculating structural prediction accuracy.

Generally, we found that all four methods performed similarly, although the dataset of RNAs is too small to be a fair comparison (Figure 4C). We also observed that v2.1 reactivities resulted in slightly higher accuracy for all four methods than v2.0 reactivities, which resulted in similar accuracies to those discussed in Loughrey *et al*. [20]. Thus all of the computational methods described in this work, coupled with SHAPE-Seq v2.1 reactivity data, can help guide researchers to more accurately model the structures of RNAs. This is particularly valuable for RNAs for which no crystal structure is available.

### 4.2 in-cell SHAPE-Seq analysis

The in-cell SHAPE-Seq technique is closely related to SHAPE-Seq v2.1, as many of the improvements in v2.1 were derived from the in-cell method [26]. The main difference, as outlined in Section 3.1, is that the RNA modification step occurs *in vivo* to provide a more natural context for RNA folding to occur. Below, we present two different examples of in-cell SHAPE-Seq data for RNAs expressed in *E. coli*.

#### 4.2.1 5S rRNA, expressed endogenously

To demonstrate how a combination of *in vitro* and *in vivo* SHAPE-Seq can provide information about how the cellular environment affects RNA folding, we used in-cell SHAPE-Seq to measure the structural characteristics of the *E. coli* 5S rRNA. We found that the reactivities we observed matched well to three-dimensional representations of the 5S rRNA from cryo-EM data fit with molecular dynamics simulations (Figure 5A) [26]. High reactivities tend to occur in unstructured loop regions, with the exception of regions that are bound by proteins within the ribosome. Regions expected to be protein-bound appear lower in reactivity, suggesting that cellular 5S rRNA is predominantly contained within the ribosome during exponential growth.

**Figure 5.**
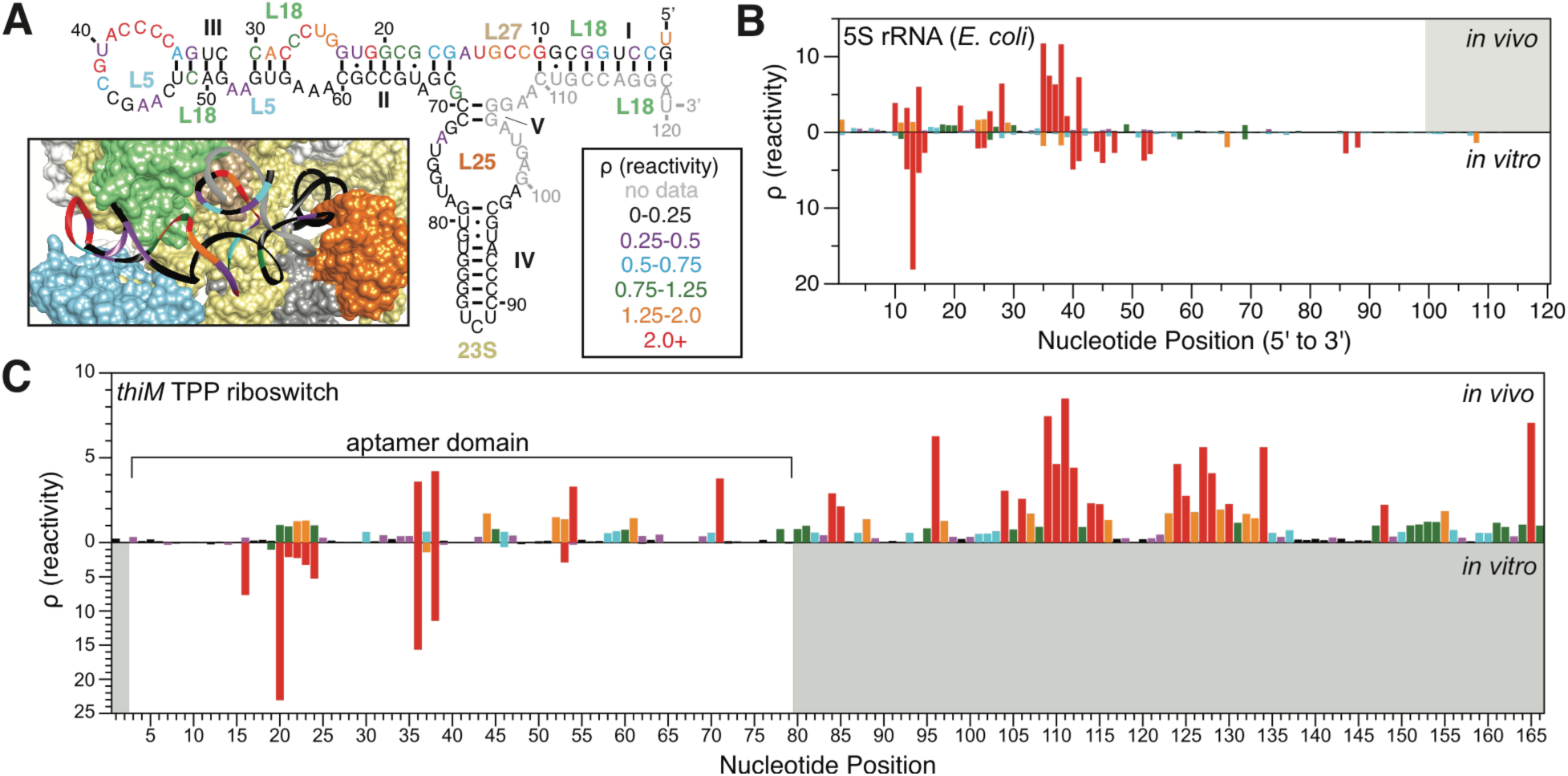
*in vitro* vs. in-cell SHAPE-Seq reactivity map comparisons for 5S rRNA and the TPP riboswitch. (A) 5S rRNA in-cell reactivities overlaid on a predicted secondary structure [71] and a three dimensional model of the 5S rRNA within the entire ribosome (inset; from PDB 4V69) [72]. Individual ribosomal proteins (L5, L18, L25, L27) and the 23S rRNA are labeled on the secondary structure near their approximate locations; helices are numbered I-V. Reproduced from Watters *et al*., 2015 [26] with permission from Oxford University Press. (B) Comparison of reactivities for the *E. coli* 5S rRNA measured in-cell (endogenous expression, top, RMDB: 5SRRNA_1M7_0007) vs. *in vitro* (bottom, RMDB: 5SRRNA_1M7_0009). Reactivities are color-coded according to (A). Clear differences in the endogenous 5S rRNA reactivities are apparent, especially for nucleotides 35–38 and 44–100, which increase and decrease, respectively relative to *in vitro*. (C) Comparison of the *E. coli thiM* TPP riboswitch measured *in vivo* (expressed from a plasmid vector, top, RMDB: TPPSC_1M7_0004) vs. an *in vitro* measurement of the adapter domain only with 5 μM TPP present (RMDB: TPPSC_1 M7_0005). Comparing the reactivities in the 5' half of the aptamer domain suggests that the riboswitch is primarily in the bound form in the cell, though differences in the 3' half suggest that the cellular environment and the aptamer sequence context affect the RNA fold. Nucleotides that were not mapped in (B) and (C) are indicated with gray.

We can also directly compare to *in vitro* 5S rRNA data to determine how the cellular environment changes the reactivity pattern (Figure 5B). For example, nucleotides 45–54 exhibit lower reactivities *in vivo*, roughly where the L5 protein is expected to bind (Figure 5B). Also, there are clusters of peaks downstream of nucleotide 54 that are near zero *in vivo*, but are highly reactive *in vitro*. Identifying these types of decreases, or increases, can reveal how different folding conditions (i.e., the cellular environment) affect RNA structure and function inside the cell.

#### 4.2.2 TPP riboswitch, expressed from a plasmid

Next, we examined the TPP riboswitch with in-cell SHAPE-Seq. Unlike 5S rRNA, the TPP riboswitch was supplied exogenously as a translational fusion with superfolder green fluorescent protein (SFGFP) from a plasmid. Interestingly, we observed a reactivity pattern in the aptamer domain that matched well to the *in vitro* reactivity map of this region in the presence of ligand (Figure 3A, 5C). This suggests that the aptamer domain is predominantly in a ligand bound confirmation in the cell. However, there are also some slight differences between the *in vivo* and *in vitro* data. For example, positions 43–46, 58–61, and 70–71 exhibit higher reactivity *in vivo* (Figure 5C). In general, it appears that the TPP aptamer domain folded *in vitro* out of context of the expression platform captures most of the interesting reactivity clusters, but potentially misrepresents some details of the riboswitch. These types of comparisons illustrate an advantage of in-cell SHAPE-Seq in that RNAs can be easily introduced with expression vectors to provide a more relevant picture of RNA folding in the cellular environment.

## 5. Experimental Considerations

### 5.1 Effect of increasing PCR cycles

In cases of low input RNA or poor cDNA yield, it may be advantageous to increase the number of PCR cycles to increase the amount of dsDNA available for sequencing. While there has been some concern that the increased number of cycles may introduce bias, we have shown in previous work that an increased number of PCR cycles does not cause substantial reactivity changes using either in-cell SHAPE-Seq (Figure 6A) or SHAPE-Seq v2.0 [20,26]. However, SHAPE-Seq v2.1 inherently requires more cycles of PCR than v2.0 to reach an equivalent library concentration because v2.1, as well as in-cell SHAPE-Seq, uses multiple forward primers that require extra PCR cycles to build the complete Illumina adapter sequences in a stepwise fashion. Therefore, to confirm that the increased number of PCR cycles in SHAPE-Seq v2.1 does not bias reactivity calculation, we sequenced our v2.1 libraries using 15 (the v2.1 standard), 18, or 20 cycles of PCR (Figure 6B-D). As expected, there was little difference in the calculated reactivities due to increased cycling for 5S rRNA, the TPP riboswitch, and adenine riboswitch.

**Figure 6.**
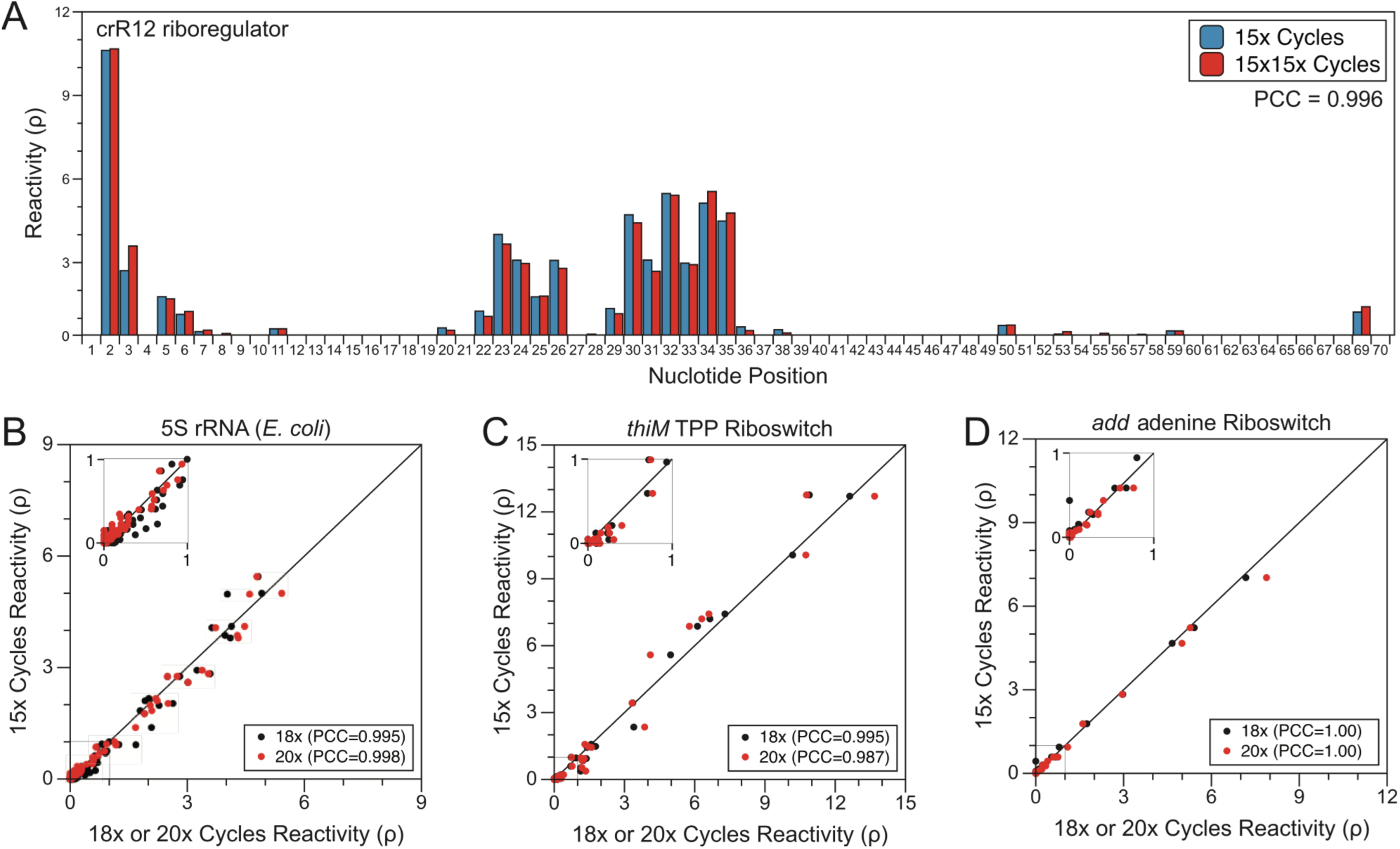
PCR does not bias SHAPE-Seq reactivity calculations. (A) Comparison of the crR12 riboregulator calculated reactivities from an in-cell SHAPE-Seq experiment [26] using either 15x cycles of PCR (15x; blue) vs. 15x cycles without PE_F, followed by another 15x cycles including PE_F (15x15x; red). A Pearson Correlation Coefficient (PCC) value of 0.996 suggests increased PCR cycling does not affect the reactivity calculation. (B) Base-wise comparison of reactivities calculated from 5S rRNA SHAPE-Seq libraries using 15x cycles of PCR vs. 18x (black) or 20x (red) cycles of PCR (RMDB: 5SRRNA_1M7_0009). A PCC near unity suggests little difference in the reactivity values calculated from libraries with increased PCR cycling. Similar analyses were done for the *thiM* TPP aptamer domain (RMDB: TPPSC_1M7_0005) (C) and *add* adenine aptamer domain (D) (RMDB: ADDSC_1M7_0008).

For experiments in which the amount of input RNA is particularly low, or improved selection against unwanted dimer side product is desired, we suggest performing the quality analysis and dsDNA library PCRs in a stepwise fashion. As shown in Watters *et al*., the amount of unwanted dimer amplification can be further reduced by first cycling with only the inner selection primer before adding PE_F for the required subsequent cycles [26]. We also found that excessive cycling with PE_F beyond 15 rounds may cause an increase in off target and dimer side product amplification, especially when the in-cell target transcript is at low abundance. Thus, for sensitive applications we recommend splitting the reaction into two phases as described above and increasing the number of cycles in the first phase if increased sensitivity is required, with the second round limited to 15 or fewer cycles after PE_F is added to complete the reaction.

Because of the ability of SHAPE-Seq v2.1 and in-cell SHAPE-Seq to selectively amplify correctly extended cDNAs, both have the capability to analyze RNAs that are present at low concentrations without altering the RNA folding conditions. To demonstrate that SHAPE-Seq v2.1 provides consistent results over a range of relevant *in vitro* RNA concentrations, we compared the reactivity maps we obtained using four different starting amounts of 5S rRNA: 1, 5, 10, and 20 pmol (Figure 7). Across all three concentrations, there is good agreement between the reactivity maps of the various starting amounts. There are several positions that exhibit slight differences, such as nucleotides 102–104, but they do not show a trend related to the starting amount of RNA. Interestingly, the nucleotides in this region exhibit fewer aligned reads relative to other nearby positions in the RNA sequence. Ultimately, it is not the amount of starting RNA, but rather the number of aligned sequencing reads used for reactivity calculation that determines data quality and consistency.

**Figure 7.**
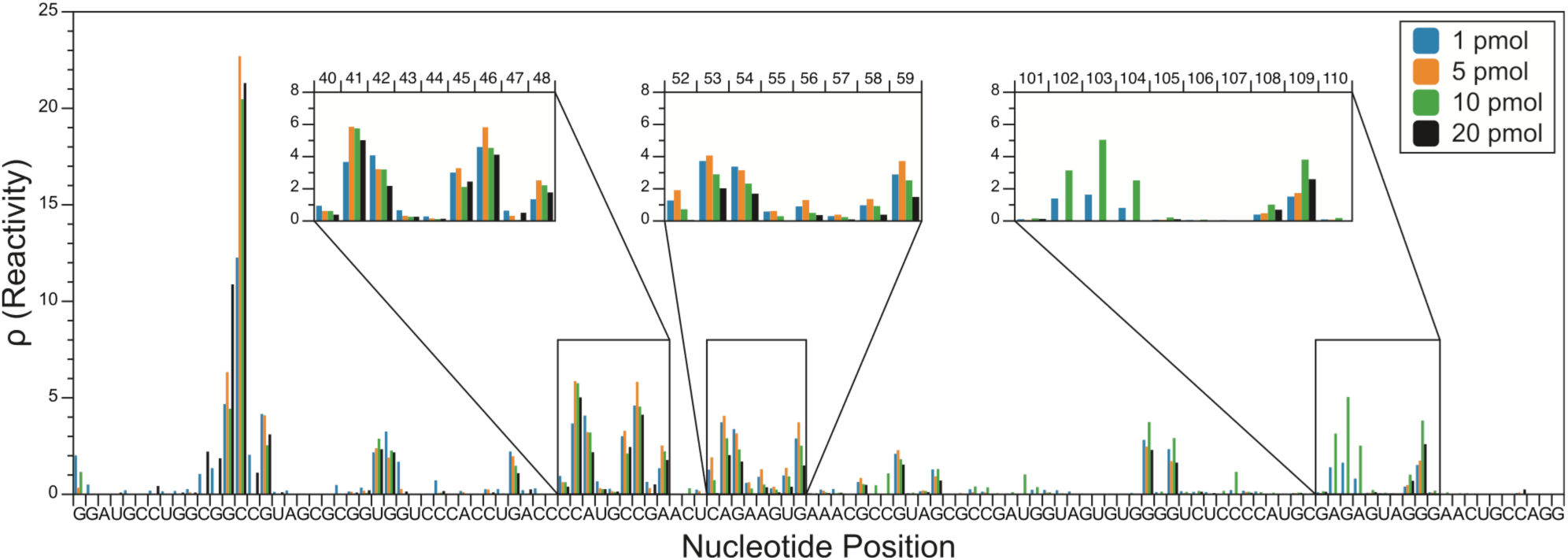
Comparison of *in vitro* folded 5S rRNA reactivity maps from different amounts of starting RNA. The same *in vitro* SHAPE-Seq experiment was performed using either 1, 5, 10, or 20 pmol of starting RNA (RMDB: 5SRRNA_1M7_0009). As expected, all of the reactivity maps are very similar, although there is some disagreement near positions 102–104 (right inset). However, these differences do not show a trend with increasing/decreasing starting RNA level and are thus likely experimental noise.

### 5.2 RT primer length and library multiplexing

One of the key features of the SHAPE-Seq improvements is the shortening of the RT primer. In its original conception, SHAPE-Seq v1.0 used long RT primers that contained the entire Illumina adapter sequence [19]. SHAPE-Seq v2.0 greatly shortened these primers and the DNA adapter sequence. It also included both TruSeq barcoding provisions and the potential for internal barcoding by using a pair (+ or -) of RT primers containing a pre-planned barcode [20]. The changes added to v2.0 improved dimer side product removal during bead purification and lowered oligonucleotide expenses.

In the current state-of-the-art SHAPE-Seq methods (v2.1 and in-cell), the RT primer is further shortened to only contain sequence that binds directly to an RNA of interest, thus requiring that all of the Illumina sequences are provided with PCR, except for those included in the DNA adapter. By adding all of the Illumina sequences this way, much more customization during library preparation for sequencing is allowed. It is also a major advantage because it decouples the preparation of the ssDNA library from the preparation of the dsDNA sequencing library. Thus, libraries can be stored for long periods of time without the concern of potential sequencing incompatibilities, as ssDNA libraries can be easily converted to dsDNA libraries at a later date with the most current barcoding and sequencing configuration. In contrast, v2.0 barcodes had to be pre-selected and could not be changed at a later date, which could cause sequencing incompatibilities between certain libraries during sequencing. Last, the shortened primers, roughly 20–35 nucleotides compared to 55–70 for v2.0 or 80–90 for v1.0, produce shorter dimer side products that are more easily removed during the bead purification steps. Thus, using short RT primers provides major advantages for SHAPE-Seq techniques. In fact, using longer RT primers with the most recent protocols will result in increased amounts of unwanted dimer ligation product, even with the improved PCR selection methods.

### 5.3 Choosing SHAPE reagents and other chemical probes

There are many choices of a chemical probe. For routine probing experiments we recommend 1M7, although there are many instances in which use of a different chemical is advantageous. The SHAPE reagents modify the 2’-OH of flexible nucleotides. Other chemicals such as DMS or CMCT directly modify the Watson-Crick face, though they are typically limited in their range of selectively. Further, the reaction time scale of the chemical probe or its ability to enter living cells may have a factor in the choice of probe to use. Below we describe some of the basic characteristics of the most common reagents to provide insight into choosing one over another for an experiment. If using a chemical other than 1M7, the reaction time and conditions may need to change relative to the described method in Section 3.1.

There are a number of SHAPE reagents that have slightly different modification properties. Three similar compounds (NMIA, 1M6, and 1M7) are based on the same anhydride scaffold and have increasingly shorter half-lives from 260 to 14 seconds for NMIA and 1M7, respectively [36,49]. Differences in the reactivities measured for the same RNA with these reagents can yield information about the ribose sugar conformational sampling based on the dynamics of modification [36]. If using NMIA or 1M6 in place of 1M7 for *in vitro* experiments, increase the reaction incubation times to 22 min or 3 min, respectively, using the same concentration as 1M7 [36].

Benzoyl cyanide (BzCN) is another type of SHAPE reagent. It reacts with a very short half-life (250 ms), reacting to completion in seconds [61]. Historically, BzCN has been used when the modification needs to take place as quickly as possible, such as with time course assays [61–63]. Because of the increased difficulty of use and elevated safety considerations required for BzCN, we recommend using 1M7 instead unless a very fast modification time is needed. When fast modifications are desired, use BzCN at 400 mM (in place of 65 mM for 1M7) and incubate the reaction for 1–2 seconds to bring it to completion.

The last major class of SHAPE reagents, consisting of NAI and FAI, have hydrolysis half-lives in the middle of the NMIA-1M7 spectrum [32]. Recently, they were further functionalized to contain an azide group that allows the addition of a biotin moiety via a ‘click’ reaction for subsequent pull-down and selection of modified RNAs only, thus reducing the required sequencing depth downstream [27]. If using NAI to modify RNA, replace the 65 mM 1M7 reagent with a 1–2 M stock of NAI or NAI-N_3_ and incubate for 15 min. Quench using a two-phase extraction (e.g. TRIzol) to remove unreacted NAI.

NAI, FAI, and 1M7, were recently shown to diffuse into living cells to modify RNAs inside the cell [26,27,31,32,64]. For in-cell SHAPE-Seq in *E. coli* we recommend 1M7 because its half-life is on a shorter time scale than cell division and RNA degradation. All three SHAPE reagents usable *in vivo* can be synthetized in one (1M7 [50], NAI, and FAI [32]) or a few steps (NAI-N_3_ [27]) from commercially available reagents. To use NAI instead of 1M7 for *in vivo* probing, replace the 13.3 μL of 250 mM 1M7 with 51.2 μL of NAI (or NAI-N_3_) 1–2 M stock solution and incubate for 15 min in place of 2–3 min before two-phase extraction to quench the reaction.

There are also a number of chemical probes that directly modify base positions. The two most popular are DMS and CMCT, which are known to preferentially modify A/C or G/U positions, respectively, although not equivalently [65]. Others, such as DEPC (diethylpyrocarbonate) and kethoxal [12], are also base specific, but are used less frequently now, mainly due to the fact that DMS and CMCT cover all four bases together and react more consistently. Unlike CMCT, DMS can enter cells to modify RNAs directly inside without forcing them to be permeable. This property and DMS’s longstanding use as a chemical probe led to its use in many of the recently published *in vivo* NGS-based probing methods [21–23].

To use DMS in place of 1M7, replace the 13.3 μL of 250 mM 1M7 with 27.75 μL of 13% DMS in ethanol (for *in vivo)* or the 1 μL of 65 mM 1M7 with 1 μL of 3.5% DMS in ethanol (for *in vitro)*, replacing the DMSO control with ethanol. Incubate for 3 minutes before quenching with 240 μL or 2.4 μL 2-mercaptoethanol for *in vivo* or *in vitro* experiments, respectively. Use two-phase extraction to purify the RNA as suggested in Section 3.1.

It should be noted that any of these chemicals should be cross-compatible with most NGS-based RNA probing methods, including SHAPE-Seq, given that most of the steps involved are for preparing the sequencing libraries. While differences in library preparation techniques do exist, as reviewed in Strobel *et al*. [11], most chemical probing methods, except for SHAPE-MaP [18], rely on the ability of modified nucleotides to block reverse transcription. Thus, by simply changing the RNA modification step SHAPE-Seq [19,20,26,52] could use DMS modification just as easily as DMS-Seq [23] could use SHAPE modification, as was done in Watters *et al*. [26].

### 5.4 Factors influencing data quality and consistency

There are a number of factors that have the potential to influence the final results that should be kept in mind while performing SHAPE-Seq experiments.

One of the biggest factors in collecting meaningful and consistent results is the importance of good RNA extractions and purifications. Poor recovery of RNA after extraction or precipitation will greatly lower the number and quality of reads aligned, mainly through increasing the amount of unwanted dimer product that is generated, as there will be less cDNA to ligate to the DNA adapter. This can be especially problematic for in-cell SHAPE-Seq during the initial total RNA extraction. We have found that extractions that become degraded, either by poor RNase-free technique or excessive delay in extracting the RNA, come out with very poor yields. Thus, careful pipetting for precipitations and extractions as well as quickly extracting total RNA, if performing in-cell SHAPE-Seq, are crucial.

A wealth of information can be gained about a library from the quality analysis steps (Figure 1). First, the rough percentage of the library that is composed of the unwanted dimer side product can be determined by observing the expected dimer peak that typically shows up around 100 nucleotides, depending on the RT primer length [26]. Second, the full-length peak can be used to ensure that the reverse transcriptase extended all the way to the 5’ end of the RNA. Also, the heights of all the peaks are indicative of the general library quality. Higher peaks suggest higher quality libraries that will need fewer cycles of PCR to prepare sequencing libraries. Last, the relative level of signal decay from reverse transcriptase stopping can be qualitatively estimated from the quality analysis and can help inform *a priori* how many reads may be required for an acceptable reactivity map.

Not surprisingly, more aligned reads generate a more accurate reactivity spectrum and reduce the run-to-run variability, or noise, between individual experiments [66]. As a rule of thumb, we suggest a minimum of 50,000 reads aligned to be confident that the maximum likelihood estimation used by Spats [33,34] generates a reliable reactivity map. However, this assumes that the reads are well distributed between the (+) and (-) samples and within the RNA, which is frequently not the case. Despite this, some RNAs actually generate reliable maps with even fewer reads, although they are RNAs that tend to have fewer, highly reactive peaks rather than large clusters of intermediate reactivities. Another rule of thumb is that roughly 10–100 reads per nucleotide position should be aligned in both channels. These values, however, represent minimums. We suggest a few hundred thousand reads to generate the cleanest reactivity maps with the least amount of variability.

The last point of consideration is the level of signal decay that occurs within the RNA. As the reverse transcriptase transcribes from the 3’ end, it has a tendency to ‘fall off’, or stop transcribing, with some probability which is increased in the (+) channel due to the presence of the chemical modifications. Correcting this signal decay is performed by the maximum likelihood model used within Spats to calculate reactivity [33,34]. However, certain nucleotide positions, either due to an inherent high ‘fall off’ rate or a high probability of chemical modification, greatly increase the signal decay rate [66]. Because Spats uses signal decay to calculate reactivity, it is resistant to errors that can occur in other analyses from rapid signal decay. However, these sharp drop-offs in read alignments can still affect Spats processing if the number of reads upstream (closer to the 5’ end) of the drop-off becomes very small. RNAs that contain these extreme drop-offs are cases were the number of reads required for a reliable reactivity map is increased. One example is highlighted above for nucleotides 102–104 in the *in vitro* 5S rRNA reactivity spectrum, which exhibits run-to-run variation and occurs in a region of fewer read alignments ( q).

### 5.5 Choosing an adapter trimming algorithm

In Loughrey *et al*., we updated the Spats data analysis pipeline to include an improved adapter trimming algorithm based on the fastx toolkit named adapter_trimmer. More recently, we have also created a version of adapter_trimmer that is based on cutadapt [67] as an alternative adapter clipping method. Further, we have relaxed the requirement that all reads aligned match perfectly to the target. The updated version of spats, as well as older versions, can be found at https://github.com/LucksLab/spats/.

### 5.6 Measures of SHAPE reactivity

The mapped read counts from the sequencing data are converted into a measure of reactivity using Spats called *θ*. Each *θ_i_* value represents the relative probability that a modification within an RNA molecule occurs at nucleotide *i*. Values for *θ* are estimated by combining a Poisson model for the number of modifications per molecule with a model of reverse transcription ‘fall off’ at the modification positions to derive a maximum likelihood estimation procedure to find the underlying *θ* that best explain the pattern of (+) and (-) read counts [33,34].

Because *θ* is a distribution describing the relative probability of modification at each position within the RNA, it is dependent on the length of the RNA according to:

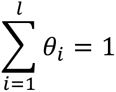

where *l* is the length of the RNA molecule. To compare RNAs of different lengths, *θ_i_* can be normalized to *ρ_i_* by multiplying by length *l* [20,26]. Using *ρ_i_* in place of *θ_i_* is also useful because it sets the reactivity values to the order of magnitude expected by secondary structure prediction algorithms as shown in Loughrey *et al*. using *m* = 1.1 and *b* = -0.3 as folding parameters [20].

## 6. Further potential improvements for SHAPE-Seq and restrained RNA folding

While we have continued to improve the SHAPE-Seq technique, there are a few areas where further improvements/extensions are still desired.

### 6.1 Going transcriptome-wide

While we generally advocate for targeted RNA structure analysis, there are many potential benefits to the recent innovations in transcriptome-wide probing techniques [22,23,25,27] that have sparked a growing interest in examining RNA structure at a global level [11]. Therefore, extending SHAPE-Seq to be able to optionally target the entire transcriptome would be valuable. To switch to total RNA structure probing, all that would be experimentally needed is an alternative RT priming step in place of the targeted approach we have chosen to follow up to this point. Random priming of total RNA is in principle easy to perform, but does not allow for selective PCR methods. Yet, it may not be required if good extension across the random primer set is achieved, leaving little to no unextended RT primer. An alternative approach is to fragment the total RNA and ligate an RT priming site, using a SHAPE-Seq v2.1 -like approach. The drawback is that the added ligation step will negatively impact library generation efficiency, and additional methods will be needed to distinguish RT fall off due to fragmentation from modification.

### 6.2 Future directions for computational folding

A common application for SHAPE-Seq reactivity data is to restrain RNA folding algorithms, as discussed extensively above. Reactivity data has been repeatedly shown to improve secondary [20,24,38,39,54] and tertiary [68] structure predictions, with secondary structure algorithms being more popular. However, there are still many cases where reactivity information alone is not enough to obtain an accurate fold. Beyond improving free energy terms and including pseudoknots, there are two main ways in which structural prediction accuracy could be greatly improved: inclusion of non-canonical base pairing and better representation of RNA structure subpopulations.

One common cause of prediction inaccuracies is the presence of non-canonical base pairing, which is pervasive in RNA structural motifs. In many cases, non-canonical base pairs are in regions that are predicted to be in single-stranded, allowing the rest of the RNA to attain a fairly accurate folding prediction. However, bases that participate in non-canonical interactions are frequently incorporated into canonical Watson-Crick pairs during computational folding, which can generate nonsensical RNA structures that are misleading for *de novo* RNA structure modeling. SHAPE reactivities often reflect non-canonical structures well if knowledge of their presence is provided *a priori* via crystal structure data, etc. Thus, even small improvements in predicting non-canonical base pairing would be of great interest to the RNA community and would greatly aid SHAPE-directed structure folding accuracy.

Another common pitfall encountered when predicting RNA structures computationally is the focus on the minimum free energy (MFE) structure. Frequently, these structures may be misleading, especially in cases where non-canonical base pairing is present, as discussed above. Further, a population of identical RNA molecules is not restrained to fold into only one structure. Rather, RNA structure is more accurately described as a combination of many structural subpopulations, which typically contain several different dominant structural motifs [69]. One method to address these subpopulations is to cluster predicted RNA structures and use these clusters to obtain a characteristic structure. Sfold and SeqFold both use this method [45,69] as well as the approach taken by Kutchko *et al*. [48]. SeqFold chooses which characteristic structure best represents the SHAPE reactivity data, but this collapses the subpopulation information into one structure. A more powerful approach would be to use SHAPE reactivity data to not only predict which structures are likely present, but also at what level they exist in a structural population. Some preliminary work has been done to understand structural populations in this manner [70] but further improvement and adoption would be beneficial to understanding potential structures when studying a new RNA.

## 7. Conclusions

SHAPE-Seq is a rapidly improving and expanding technique for characterizing the RNA structure-function relationship both *in vitro* and *in vivo*. In this work, we have presented different experimental approaches for characterizing RNA structures both *in vitro* and *in vivo* and showed how to use the structural information obtained to computationally predict what RNA structures were present in the experimental conditions. As our data suggest, SHAPE-Seq is a robust technique that has been updated to be simpler to perform through the use of selective PCR and optimized library construction steps. Further, the wide variety of computational tools available for RNA secondary structure prediction can be used to help interpret SHAPE-Seq results, with many of them able to directly incorporate reactivity data to improve structure prediction accuracy. We believe that the widespread adoption of SHAPE-Seq methods backed by computational tools will continue to drive the discovery of new insights in RNA structural biology.

## Acknowledgements

This material is based upon work supported by the Tri-Institutional Training Program in Computational Biology and Medicine [NIH training grant T32GM083937 to AMY] and a New Innovator Award through the National Institute of General Medical Sciences of the National Institutes of Health [grant number 1DP2GM110838 to J. B. L.]. K.E.W. is a Fleming Scholar in the School of Chemical and Biomolecular Engineering at Cornell University. The content is solely the responsibility of the authors and does not necessarily represent the official views of the National Institutes of Health.

